# Differences in activity and stability drive transposable element variation in tropical and temperate maize

**DOI:** 10.1101/2022.10.09.511471

**Authors:** Shujun Ou, Tyler Collins, Yinjie Qiu, Arun S. Seetharam, Claire C. Menard, Nancy Manchanda, Jonathan I. Gent, Michael C. Schatz, Sarah N. Anderson, Matthew B. Hufford, Candice N. Hirsch

## Abstract

Much of the profound interspecific variation in genome content has been attributed to transposable elements (TEs). To explore the extent of TE variation within species, we developed an optimized open-source algorithm, panEDTA, to *de novo* annotate TEs in a pan-genome context. We then generated a unified TE annotation for a maize pan-genome derived from 26 reference-quality genomes, which revealed an excess of 35.1 Mb of TE sequences per genome in tropical maize relative to temperate maize. A small number (n = 216) of TE families, mainly LTR retrotransposons, drive these differences. Evidence from the methylome, transcriptome, LTR age distribution, and LTR insertional polymorphisms revealed that 64.7% of the variability was contributed by LTR families that were young, less methylated, and more expressed in tropical maize, while 18.5% was driven by LTR families with removal or loss in temperate maize. This study demonstrates the use of a comprehensive pan-TE annotation to reveal the driving role of TEs in within-species genomic variation via their ongoing amplification and purging.

## Main

Eukaryotic genomes are largely comprised of transposable elements (TEs). For example, 54% of the completed human genome consists of repetitive sequences^1^. In maize, where TEs were first discovered, 85% of the genome consists of TE sequences, of which 75% are long terminal repeat (LTR) retrotransposons^2^. In many species, TEs are intertwined with genes and, as a result, can have functional consequences by altering transcript structure or regulation^3^. The important maize *bz* locus varies from 50 - 160 kb among different genotypes due to variable TE insertions and deletions^4^. TEs can also alter transcript abundance^3^ as seen with a TE insertion upstream of the *tb1* gene^5,6^ that results in increased expression, which enhances apical dominance and reduces tillering in domesticated maize. Likewise, TE insertions in the promoter regions of *ZmCCT9* and *ZmCCT10* (ref ^7,8^) alter gene expression leading to early flowering and long-day adaptation in temperate maize. Expression differences can also result from insertions into intron sequences, such as a Mutator-like TE in an intron of *DSX2* (ref ^9^) that enhances expression of the gene leading to carotenoid accumulation and yellow kernels. Total TE content in genomes has also been linked to environmental gradients such as altitude in maize^10^ and biotic and abiotic responsiveness in tomato^11^, suggesting an adaptive role for TEs.

Despite the importance of TEs in genome evolution and crop improvement, there have been a limited number of genome-wide studies of TE variation within a species. Instead, the vast majority of studies characterizing TE content have been in the context of a single reference genome^11–13^ due to previous challenges of assembling genomic regions containing these highly repetitive elements, a limited number of genome assemblies within species, and challenges with genome-wide annotation of the different classes of TEs^14^. With the continued advancements in long-read sequencing technologies and improved assembly algorithms, there is a growing movement in genomics towards pan-genomics-based approaches^3,15,16^. The recent publication of 26 reference-quality genome assemblies for the founders of the maize nested association mapping (NAM) population^17^ provides an unprecedented opportunity to explore intraspecies-level variation in TE content and to directly test the relative contribution of mechanisms that may underlie variation in the abundance of individual TE families. These genomes all have gold-quality assemblies based on the LTR Assembly Index^17,18^, and, as such, any observed differences in TE content likely reflect the biology of these genomes rather than assembly artifacts. The panel also includes an even balance of tropical- and temperate-adapted germplasm that exhibits marked differences in flowering time, disease resistance, plant height, and other important agronomic traits that may be driven by variation in TE content (reviewed in ref ^19^).

To study variation in TE content across these 26 maize genomes, we developed panEDTA based on the Extensive *de-novo* TE Annotator (EDTA) pipeline^14^, facilitating structural and homology-based *de novo* TE annotation in a pan-genome context. panEDTA first uses EDTA to identify structurally intact TEs in each genome, then combines *de novo* libraries into a comprehensive pan-genome library, which is used to consistently reannotate each genome, allowing fragmented elements in a genome to be homology annotated, and provide uniform TE family names across all family members in all genomes (**Fig. 1a**). The original EDTA pipeline and the panEDTA modules are available for download at (https://github.com/oushujun/EDTA) and have both been implemented in the browser-accessible cloud-based platform Galaxy^20^.

**Figure 1.**
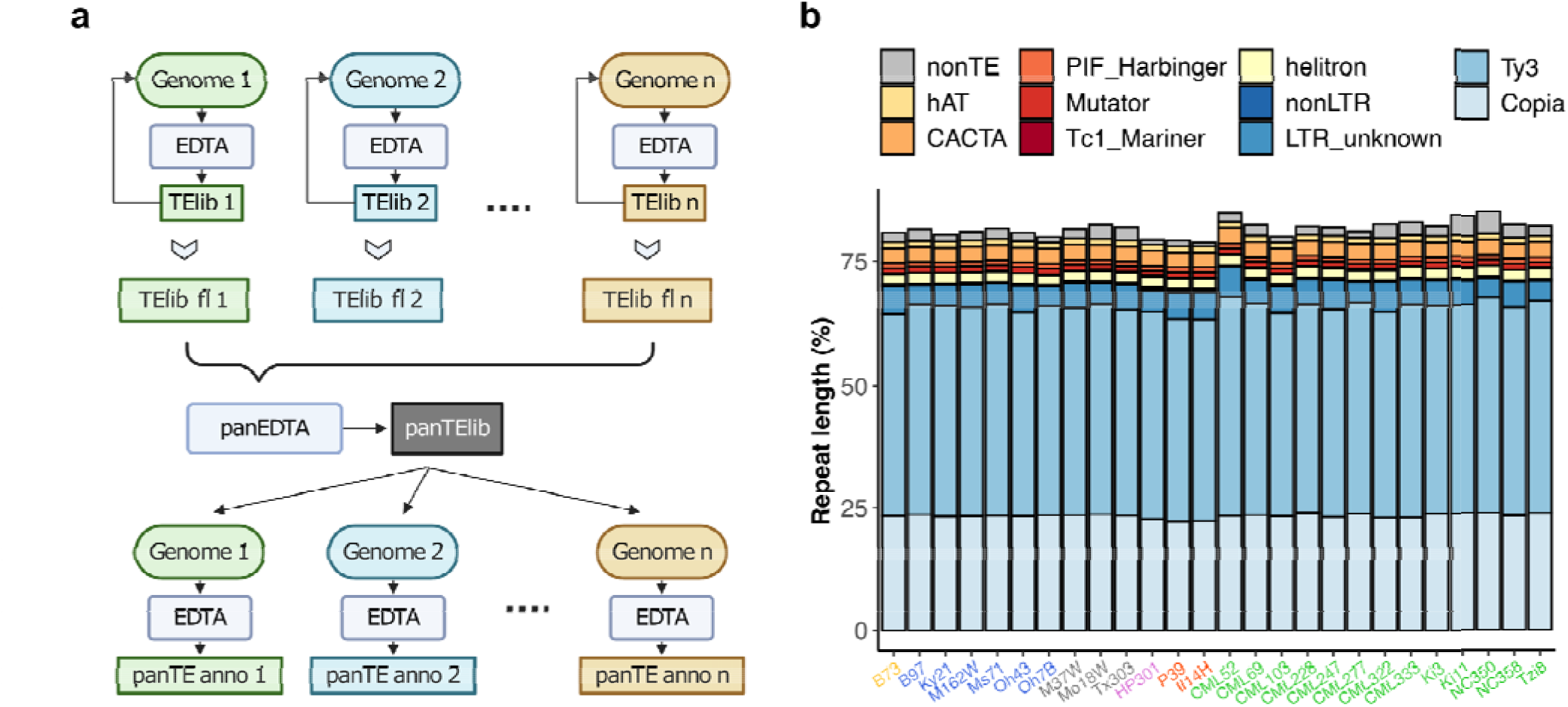
Pan-genome annotation of 26 maize NAM founders using panEDTA. **a**. The panEDTA workflow. The EDTA pipeline is used to annotate each genome independently, and the resulting individual TE libraries are filtered based on copy number and combined to form a non-redundant pan-TE library, which is used to reannotate each genome for a consistent pan-genome TE annotation. **b**. panEDTA annotation of 26 maize NAM founders. Maize lines were grouped into stiff-stalk (yellow), non-stiff-stalk (dark blue), popcorn (pink), sweet corn (red), admixed maize (gray), and tropical maize (green).

Using panEDTA, we annotated the 26 NAM founder assemblies and identified 17,473 pan-genome TE families and 269,847 unclassified low-copy TEs (**Fig. 2a, Suppl. Fig. 1a**). Together, TEs and non-TE repeats contribute an average of 88.2% of the NAM founder genomes (**Fig. 1b)**, consistent with previous reports in maize^2,17^. panEDTA in maize, as well as in rice and Arabidopsis (**Suppl. Fig. 2**; **Suppl. Fig. 3a)**, annotated a similar number of total bases of TE sequence but with a substantial improvement in the consistency of element classification (**Suppl. Fig. 3b-d**). The majority of maize TE families are small, with 89.7% of pan-genome families comprising less than 100 kb per genome (**Suppl. Fig. 4**). Collectively, these small families comprise only 6.6% (SD = 0.1%) of total TE content (**Fig. 2a**). In contrast, the 1,805 largest families contribute more than 90% (SD = 1.2%) of total TE content per genome (**Fig. 2a, Suppl. Fig. 4**), with the 50 largest TE families contributing 52% (**Fig. 2b**). Many (72.3%) TE families, including all families larger than 100 kb (**Suppl. Fig. 1bc**), occur in all 26 NAM founder lines (**Suppl. Fig. 5**), and 91% of pan-genome TE families are found when sampling as few as four lines (**Fig. 2c**).

**Figure 2.**
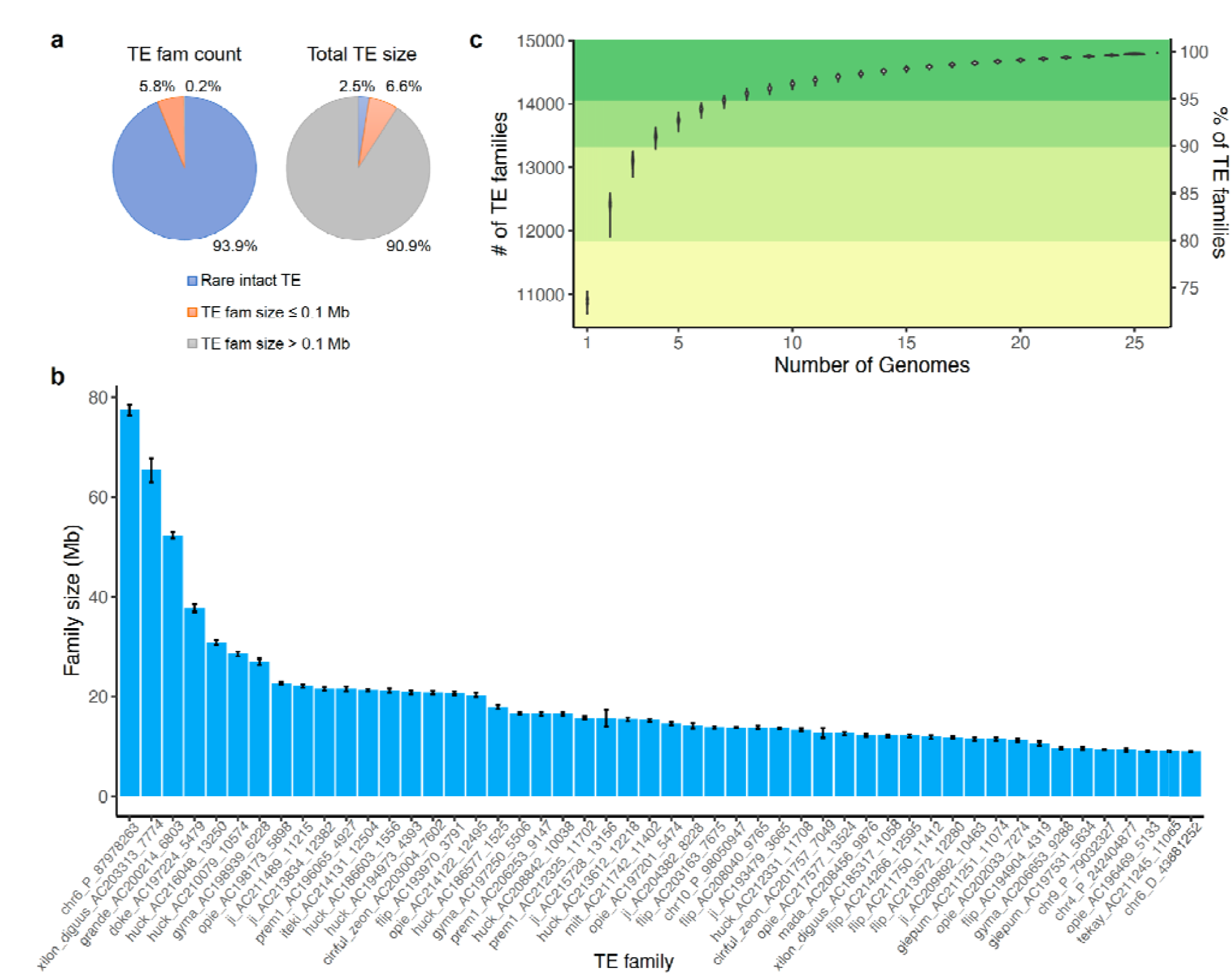
The landscape of transposable elements in the maize NAM founder genomes. **a**. Pan-genome TE family number and size. Rare intact TEs are those not classified by the 80-80-80 rule and are mostly single-copy elements. **b**. Mean size of the 50 largest TE families in the NAM founder genomes. All these families are LTR retrotransposons. The error bars denote the standard deviation among the NAM founder genomes. **c**. Summary of the number and percentage of pan-genome TE families in the NAM founder genomes. The order of genomes was shuffled 1,000 times.

While many families are consistently observed across genomes, their abundance varies. Principal component analysis (PCA) based on family size across genomes revealed substantial divergence between temperate and tropical maize along the first PC (**Fig. 3a**), a finding consistent with population structure based on SNP-based PCA and phylogenetic analysis (**Suppl. Fig. 6**). Divergence between tropical and temperate genomes based on TE family size was driven by a small number of highly variable families, with only 216 families exhibiting per genome differences greater than 0.025 Mb (**Fig. 3b)**. The ten families that varied most in size (**Fig. 3c**) were also among the 50 largest TE families in maize genomes (**Fig. 2b**). These families were significantly larger in tropical than temperate genomes (*t*-test, *P* < 1.0e-10; **Fig. 3c**), with admixed genotypes having intermediate family sizes. Those families that were larger in tropical genomes contained a combined average of 51.9 Mb more TE sequence per genome in tropical lines, while those larger in temperate lines contained an average of 16.8 Mb more sequence, resulting in a net difference of 35.1 Mb between tropical and temperate genomes. Structurally intact TEs contributed 50.8% (17.8 Mb) of the total TE variation, which was significantly more than the whole-genome average of 30.7% structurally intact TEs (Fisher’s exact test, *P* = 0.02; **Suppl. Fig. 7a**). LTR retrotransposons contributed 98.1% (34.4 Mb) of the total TE variation, with *Ty3* elements showing the largest size difference (21.5 extra Mb in tropical genomes; **Fig. 3d**). The remaining TE variation between tropical and temperate genomes was contributed by terminal inverted repeat (TIR) transposons (4.4%, with CACTA contributing 3.3%), Helitrons (-2.6%), and long interspersed nuclear elements (LINEs; -0.02%; **Fig. 3d**).

**Figure 3.**
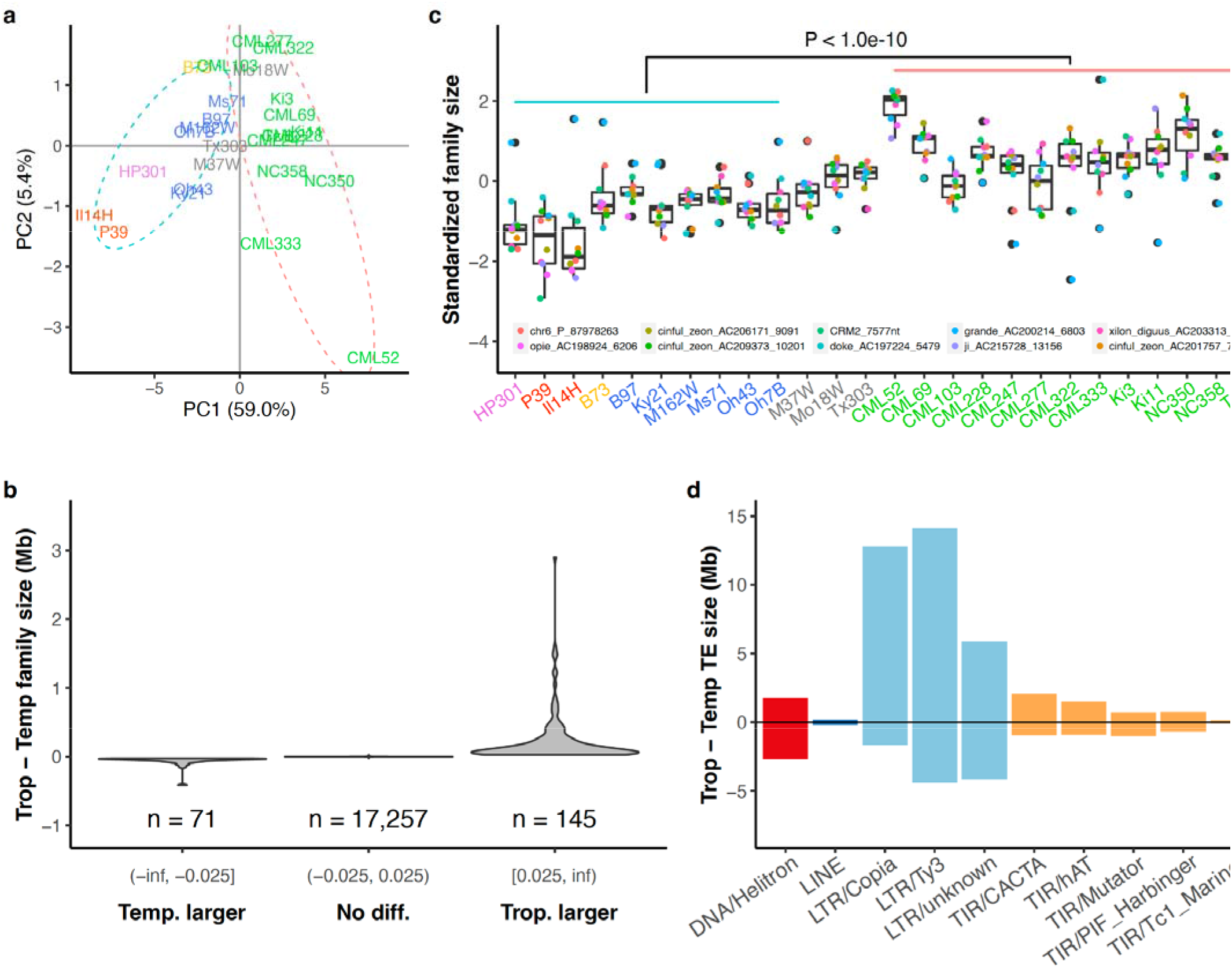
Family size variation between tropical and temperate maize genomes. **a**. Principal component analysis based on pan-TE family size in the NAM founder genomes. A total of 17,473 families were included, and the size of the family was determined by the number of base pairs in each genome. Dashed ellipses indicate tropical (pink) and temperate (blue) genomes. **b**. Distribution of TE family size difference between tropical and temperate lines. Families are divided into three categories with a cutoff of +/- 0.025 Mb difference. **c**. Distribution of the top 10 TE families with the greatest size variation among the NAM founder genomes, which are all LTR families. The size of each family was standardized to have mean = 0 and standard deviation = 1 within NAM founder lines. Maize lines were grouped into temperate maize [popcorn (pink), sweet corn (red), stiff-stalk (yellow), non-stiff-stalk (dark blue)] as indicated by the blue line on top of the boxes, admixed maize (gray), and tropical maize (green, as indicated by the pink line on top of the boxes). The box shows the median, upper, and lower quartiles. Whiskers indicate values□≤□1.5× interquartile range. Black dots indicate outliers. **d**. TE family size difference between tropical and temperate lines in TE superfamilies. Positive values represent families that are larger in tropical genomes and negative values represent families that are larger in temperate genomes.

LTR family size can be increased or reduced through retrotransposition or illegitimate recombination, respectively. LTR sequences proliferate via reverse transcription^21^, and more active proliferation could increase relative family size in some genomes. Alternatively, intact LTR elements can be reduced to solo LTRs via illegitimate recombination^22^, and preferential removal could decrease relative family size. Consequently, the solo:intact ratio of each family reflects the extent of LTR removal via illegitimate recombination, with a high solo:intact ratio suggesting substantial removal^22^. We evaluated the contribution of proliferation and removal to LTR family size variation between tropical and temperate genomes by comparing both family size and the solo:intact ratio of each LTR family. Amplifying families were defined as families having non-significant differences in removal intensity (*i*.*e*., solo:intact ratio ≈ 1) between tropical and temperate genomes, but significant differences (*t*-test, *P* ≤ 0.05) in family size (**Fig. 4a**). Amplifying families were observed in both tropical (n = 145) and temperate (n = 59) genomes, with tropical amplifying families contributing 19.2 Mb of additional sequences on average in tropical genomes and temperate amplifying families contributing an average of only 0.5 Mb of additional sequence in temperate genomes (net of 18.7 Mb, 53.3% of total differences; **Fig. 4bc, Suppl. Fig. 8ab**). Families undergoing removal were then identified as those with significantly different removal intensities and significantly different family sizes (**Fig. 4a**). These families net an extra 10.5 Mb (29.9% of total) of LTR sequences in tropical genomes (**Fig. 4bc, Suppl. Fig. 8ab**), of which 11.0 Mb is the product of stronger removal in temperate genomes. Compared to intact TEs in tropical amplification families, those in temperate removal families have shorter LTRs (median 1,226 bp vs. 1,310 bp; **Suppl. Fig. 9a**), fewer coding domains (30.4% vs. 65.0% percent of elements having complete sets of coding domains; **Suppl. Fig. 9cd**), and shorter overall element length (median 7,571 bp vs. 9,478 bp; **Suppl. Fig. 9b**). Overall, the emerging picture is that genome size variability between tropical and temperate maize genomes has largely been driven by TE families that are categorized as tropical amplification, with a lesser contribution from elevated LTR removal in temperate genomes.

**Figure 4.**
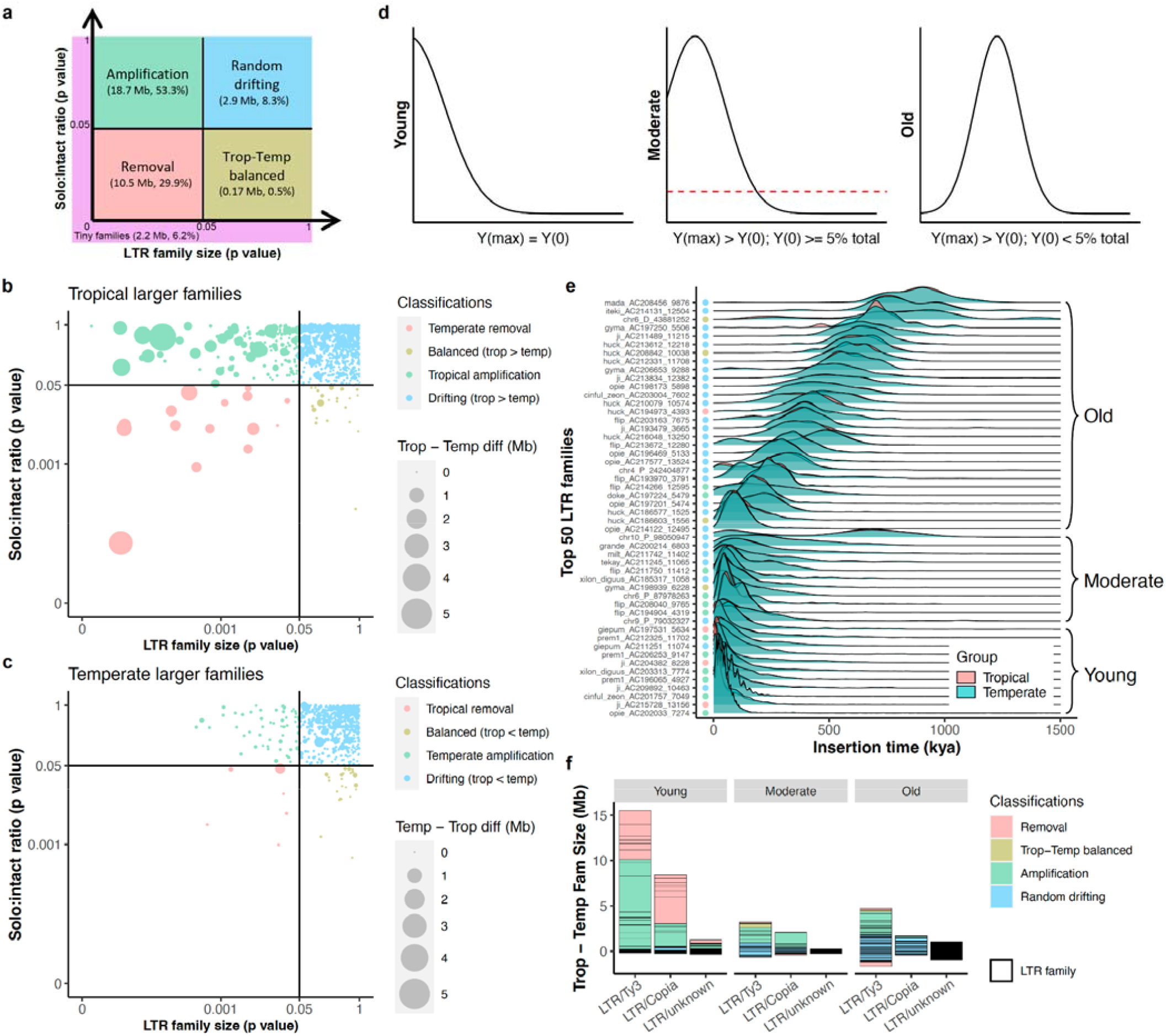
Contribution of LTR amplification and removal to genome size differentiation. **a**. Classification schematic for LTR families based on solo:intact ratio and size differentiation between tropical and temperate genomes. Numbers in Mb indicate cumulative differences in family sizes between tropical and temperate genomes and their contribution to total TE differences. **b**. LTR family classification for families that are larger in tropical genomes. **c**. LTR family classification for families that are larger in temperate genomes. In both **b** and **c**, each dot represents an LTR family, and the size of each dot scales to the absolute family size difference, and x and y axes were log10 scaled. **d**. Classification schematic for age of LTR families based on the peak frequency of insertion time. **e**. Age landscapes of the 50 largest LTR families in tropical (pink) and temperate (blue) maize genomes with overlaps shown in green. Dots indicate family classifications using the coloring scheme shown in **a. f**. The accumulated TE size differentiation contributed by different LTR superfamilies (*Ty3, Copia*, and unknown) in different age groups (Young, Moderate, and Old). Each box represents the contribution of an LTR family.

Differences in the abundance of TE families between tropical and temperate maize could have occurred at any time since the divergence of these two groups. The level of activity over time for a family can be monitored by the age distribution of individual elements within the family, which is determined based on sequence divergence between the two terminal regions of an LTR element^23,24^. We therefore next evaluated if the per family age of LTRs varied between tropical and temperate genomes along with their contribution to TE content variation between these groups. Families were classified as “Young” when the age of elements in the family peaked at 0 million years (MY; no intra-element divergence between LTRs; **Fig. 4de**). “Moderate-aged” families also have an appreciable fraction (> 5%) of 0-MY aged elements, but with peak activity in the past. Finally, families were classified as “Old” if they contained few or no (≤ 5%) 0-MY-aged elements (**Fig. 4de, Suppl. Table 1**). Overall, Young LTR families contributed 69.8% of the TE-content difference between tropical and temperate genomes, which is significantly more than the expected genomic abundance of 20.2% (Fisher’s exact test, *P* = 2.7e-5). Of these, Young *Ty3* LTRs preferentially amplified in tropical genomes and contributed 9.9 Mb (28.2%) of the TE content variation genome-wide (**Fig. 4f**), with the *xilon_diguus_AC203313_7774* family (the second largest of all TE families; **Fig. 2b**) contributing 3.9 Mb of the size difference. Conversely, Young *Copia* LTRs that were preferentially removed in temperate genomes contributed 4.9 Mb or 14.1% to TE-content difference (**Fig. 4f**), with the *ji_AC215728_13156* family contributing 2.9 Mb of this size difference. Interestingly, the *xilon_diguus* family is found in DNA regions that coincide with late DNA replication during the S phase, while the *ji* family is found in early replication regions that are usually enriched with genes^25^. Together, Young tropical amplifying *Ty3* and Young temperate removing *Copia* LTR families contributed nearly half (42.3%) of the observed TE-content difference between tropical and temperate genomes, while only making up an average of 11.8% of all TE content in the genome. For structurally intact LTR elements, Young families contributed 91.5% of the intact LTR size differentiation (**Suppl. Fig. 7b**), an over-representation compared to the expectation of 33.1% from intact Young LTR families (Fisher’s exact test, *P* < 1.0e-10; **Suppl. Fig. 7c**), suggesting structurally intact elements from Young families are driving much of the differences between tropical and temperate genomes. However, Old LTR families, especially *Ty3* LTRs, also contributed to substantial TE size differentiation (8.9%) between tropical and temperate genomes (**Fig. 4f**). The removal extent (*i*.*e*., cumulative solo:intact ratio) of all Old families was six to ten times higher than all Moderate and Young families, respectively (**Suppl. Fig. 8cd**).

Activity (*i*.*e*., transcription and amplification) of LTR retrotransposons often requires suppression of methylation^21,26–30^. The lack of methylation in the CHG context, where H = A, C, or T, is particularly informative in defining euchromatic regions. Such CHG UnMethylated Regions (UMRs) that originate within 5’ LTRs (UM-5’LTRs) rather than internal sequences of the element (**Suppl. Fig. 10**) are more likely to lead to transcription of the full-length TE. From previously reported genome-wide UMRs^17^, we identified a total of 14,074 that were UM-5’LTRs across the 26 genomes (**Suppl. Table 2**). These UM-5’LTRs were significantly enriched in tropical amplification families (observed 53.5% compared to the expected 38.7%, Fisher’s exact test, *P* < 1.0e-10) and significantly depleted in temperate removal families (observed 7.7% compared to the expected 13.9%, Fisher’s exact test, *P* < 1.0e-10; **Fig. 4f, Fig. 5a**). The fraction of UM-5’LTRs in tropical amplification families is 2.4 times that observed in temperate removal families (1.53% vs 0.63%). These UMRs rarely extended into the TE coding regions (**Suppl. Fig. 10**), and their average length ranged from 375 - 550 bp for *Ty3* and *Copia* elements (**Suppl. Fig. 11**). UMRs of this length will span 2 - 3 nucleosomes (nucleosome repeat length is ∼190 bp in maize^31^), which may allow initiation of transcription. Notably, Young *Ty3* UM-5’LTRs are 1.2 times more frequent in tropical than temperate genomes on average (Fisher’s exact test, *P* = 0.04; **Fig. 5a**), suggesting higher transcription potential of these LTR elements in tropical genomes.

**Figure 5.**
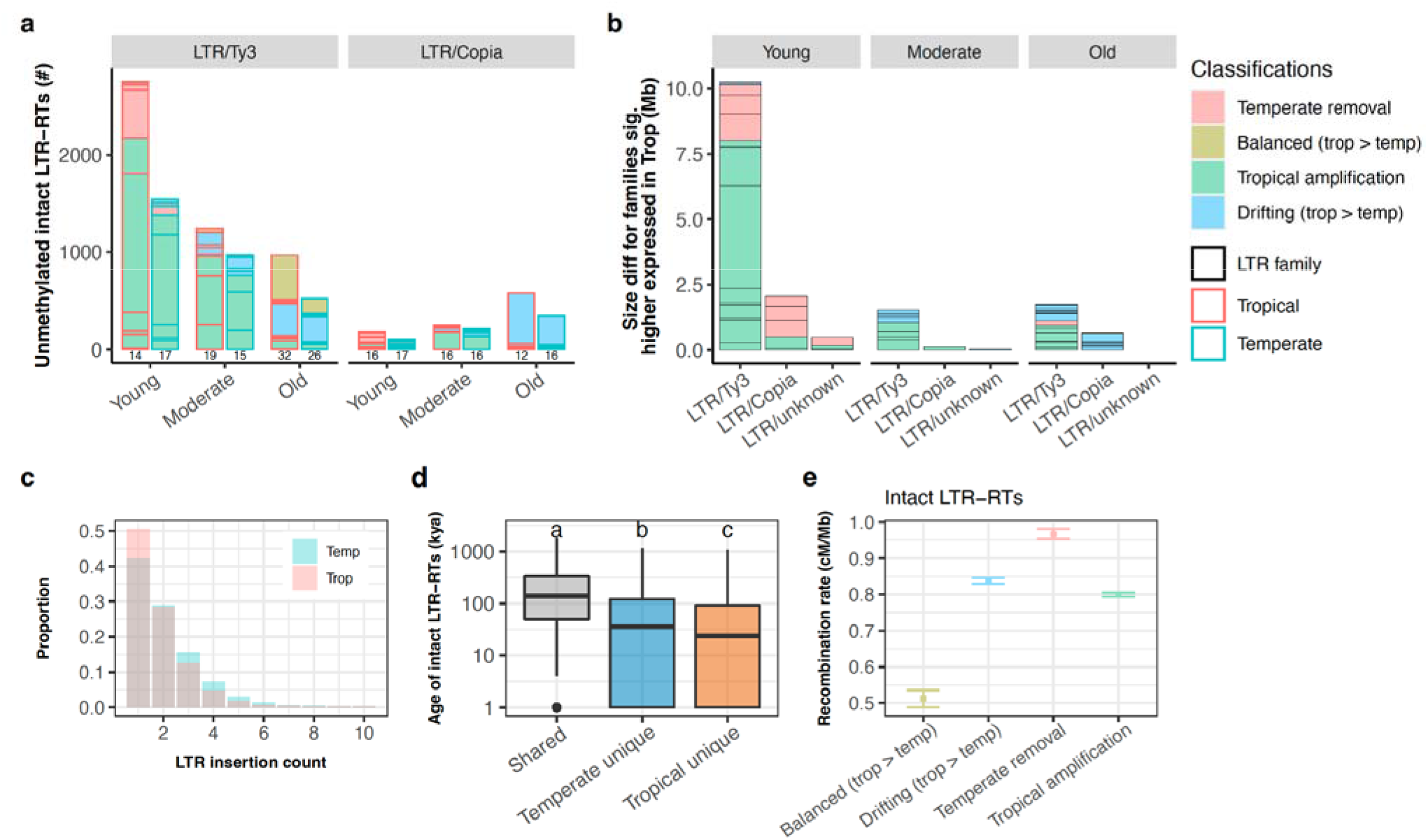
Molecular characterization of LTR families in maize. **a**. The number of intact LTR retrotransposons (LTR-RTs) carrying unmethylated regions. Data from tropical and temperate genomes are shown in side-by-side red and blue boxes, respectively. The number of families represented is indicated below each column. **b**. The accumulated family size difference between tropical and temperate genome for LTR families expressed significantly higher in at least one tissue (and with consistent directionality in all tissues with expression) in tropical genomes. **a, b**. The size of each box represents the number of LTR elements or effect size of each family, and only families that are larger in tropical genomes are shown. **c**. LTR insertion frequency spectrum in tropical (pink) and temperate (blue) genomes. Only sites younger than 20 kya were kept to increase accuracy of the polarization of the spectrum. No missing data filter was applied. **d**. The age of intact LTR elements that were shared or unique in tropical and temperate genomes. The y-axis was log10 scaled. Different letters indicate significant differences in age (Tukey’s HSD, *P* < 0.05). The box shows the median, upper, and lower quartiles. Whiskers indicate the 1.5× interquartile range. Black dots indicate outliers. **e**. Mean recombination rate for genomic neighborhoods of all intact LTR-RTs. Error bars indicate 95% confidence interval estimated from 1,000 times of bootstrap resampling.

To evaluate the functional impact of these unmethylated LTRs on transcriptional activity, we quantified TE transcript abundance across 10 diverse tissues throughout development that were previously sequenced^17^. The repetitive nature of transposable elements makes quantification of transcript abundance challenging on a per-element basis. We therefore evaluated transcript abundance on a per-family basis within each genome as previously described^32^, and conducted differential expression analysis between tropical and temperate genomes (**Suppl. Fig. 12ab**). The total transcript abundance of each TE family is positively correlated to the size of each family (Pearson’s r = 0.44, *P* < 1.0e-10; **Suppl. Fig. 13**). Particularly, the total abundance of tropical amplification families was 20.8 times higher than that of temperate removal families (**Suppl. Fig. 12c**). When normalized with the total sequence length, the total abundance of tropical amplification families was still 4.8 times higher than that of temperate removal families (**Suppl. Fig. 12de**), suggesting more active transcription of tropical amplification families. We identified 1,581 LTR families that had consistently higher abundance in tropical genotypes across all tissues (Wald test, FDR < 0.05), which explained 59.5% of TE family size differences between tropical and temperate genomes. Among these 1,581 LTR families, tropical amplification families contributed a significantly larger portion (5.7 times) than the random expectation (Fisher’s exact test, *P* = 0.048), collectively explaining 32.0% of TE family size differences between tropical and temperate genomes (**Fig. 5b, Suppl. Fig. 14**). In contrast, the contribution from temperate removal families did not exceed random expectations (Fisher’s exact test, *P* = 0.34). All but one LTR family (144/145) that possessed at least one UM-5’LTR were differentially expressed between tropical and temperate lines (**Suppl. Fig. 15a**). A total of 16 tropical amplification families that had significantly and consistently higher abundance in tropical genomes also possessed at least one UM-5’LTR in a tropical genome (FDR < 0.05). These 16 families contributed 25.7% of LTR size differences between tropical and temperate genomes (**Suppl. Fig. 15b**). Taken together, these results show accumulation of LTR sequences in tropical maize genomes is associated with lack of methylation and increased expression of tropical amplification families in tropical maize genomes.

Patterns of abundance, dating of TE activity, methylation, and expression results all suggest TE content in tropical and temperate genomes has recently evolved. To further assess this possibility, we evaluated population genetic evidence for more recent TE activity in these genomes. We identified syntenic LTR (synLTR) loci between pairs of genomes and then summarized across pairs to obtaining insertion frequencies of individual TEs at the population level (**Suppl. Fig. 16, Suppl. Fig. 17a-d**). As expected under a model of recent transposition, most LTR insertions were rare (**Suppl. Fig. 17ef**). Additionally, LTR insertion frequency was positively correlated with age (Pearson’s r = 0.89, *P* < 0.0005) and distance from genes (Pearson’s r = 0.90, *P* < 0.0004, **Suppl. Fig. 17gh**). Since the majority of variation in genome-wide LTR content was driven by tropical amplification families (**Fig. 4a**), we expected and observed excess of rare (Frequency < 20%) LTR insertions in tropical genomes (Fisher’s Exact Test, *P* < 1.0e-10; **Fig. 5c, Suppl. Fig. 18a**). This trend is also consistent with a loss of rare variants in temperate lines due to their demographic bottleneck (**Suppl. Fig. 18bc**; ref ^33^), an alternative explanation of these findings. To further assess the origin of this result, we identified LTR insertions that were unique in either the tropical (n = 7,790) or temperate (n = 5,188) groups. The age of unique LTR insertions in tropical genomes was significantly younger than unique insertions in temperate genomes (Tukey’s HSD Test, *P* < 1.0e-10, **Fig. 5d**), suggesting more recent amplification activity in tropical genomes.

Finally, to explore a mechanism for temperate removal of TEs, we considered that recombination rate might influence removal frequency and the prevalence of solo LTRs^34^. We estimated the meiotic recombination rate for genomic neighborhoods of each intact LTR element and each solo LTR using a composite recombination map from recombinant inbred lines (RILs) derived from the NAM founder parents^35,36^. Overall, temperate removal families were located in genomic regions with a significantly higher recombination rate compared to tropical amplification families across all of the genomes (Pairwise Permutation Test, *P* < 0.05; **Fig. 5e)**. As previously reported, illegitimate recombination is the dominating force in counteracting LTR amplification^22,37–39^, and the higher level of recombination in regions that contain temperate removal families suggests that recombination is, to some extent, driving the variation in size of these families between tropical and temperate genomes.

In summary, panEDTA allowed, for the first time, access to high-quality pan-genome TE annotations in maize, and characterization of the previously undescribed TE content variability between tropical and temperate maize genomes. We showed that the larger TE content in tropical genomes was mainly due to two mechanisms: an excess in LTR proliferation in tropical genomes and elevated LTR removal in temperate genomes. By employing multidimensional data including methylome, transcriptome, and pan-genome LTR polymorphisms information, we found that LTR families proliferating in tropical lines were less methylated and more actively transcribed. However, characterizing evolutionary dynamics and molecular features of TEs are still challenging due to TEs’ repetitiveness and the lack of tools suitable to their individual analysis, which is a major task for future development in computational biology that will be facilitated by continued improvements in long-read sequencing.

Temperate maize has been subject to population bottlenecks and inbreeding^40,41^, and thus harbors less genetic variation compared to tropical maize. Furthermore, maize genome size is likely an adaptive trait that negatively correlates with flowering time for latitudinal and elevational adaptation^10,42,43^. We observed reduction in TE family sizes in temperate maize that may not be solely explained by recent demography. Likewise, both selection and population expansion may have contributed to the excess of rare LTRs in tropical genomes. The more abundant LTRs of tropical genomes may contribute to increased allelic and functional variation that support robust tropical populations in the face of diverse biotic and abiotic challenges^42^. A better understanding of LTR activation and purging will provide insights, not only into the genetic and genomic dynamics in maize, but also into the general contribution of LTR retrotransposons to biodiversity in plants. The selective determinants of TE content are likely diverse, occurring at the TE family level based on insertional preference (*e*.*g*., genic vs. non-genic regions; ref ^44^) or at the genome level due to factors including environmental stress and demographic change.

## Online Methods

### Development of panEDTA

We developed panEDTA to optimize pan-genome TE annotation, which was incorporated into the current EDTA version (v2.1). Briefly, panEDTA initially annotates each genome individually for structural and homology annotation of TEs, then identifies and retains exemplar sequences with at least three full-length copies in a single genome. The previously proposed 80-80-80 rule is applied to determine full-length copies that meet these criteria, which requires ≥80% of the TE covered by sequences with ≥80% identity and with a minimal length of 80 bp^45^. By eliminating low-copy and incomplete sequences in individual libraries, panEDTA is able to filter out a large portion of potentially false TEs that will aggregate when multiple genomes are jointly annotated. Removing low-copy exemplars also keeps the pan-genome library in a reasonably small size for computational efficiency, while the potential loss of sensitivity due to the removal of low-copy exemplars in a single genome is offset by the compilation of multiple TE libraries into the final pan-genome filtered library. Sequence redundancy of the pan-genome library is removed using the 80-80-80 rule, and the remaining sequences contain a single exemplar sequence for TE families across the pan-genome. Finally, this filtered pan-genome library is used to re-annotate all genomes in a pan-annotation, including both structurally intact and fragmented TEs, with consistent family IDs across genomes. Structurally intact TEs that were not able to be classified into a family by the 80-80-80 rule are named by their genome coordinates and regarded as rare intact TEs.

We have also integrated panEDTA and the original EDTA software into Galaxy^20^, a browser-accessible cloud-based workbench for scientific computing. In brief, the Galaxy instance of panEDTA utilizes the publicly available panEDTA BioContainer and several modified scripts to render panEDTA with a graphical user interface. This implementation of panEDTA maintains the previous functionality of EDTA and first identifies structurally intact TEs in user-selected genomes. It then combines these genomes into a comprehensive library using the panEDTA algorithm and then reannotates the user-selected genomes and provides uniform TE family names across all family members in all genomes. Additionally, the Galaxy instance of panEDTA has been parallelized to run multiple instances of EDTA to decrease initial genome annotation time. We also optimized the execution of EDTA to detect Helitrons, LTRs, and TIRs in separate and simultaneous instances and then combine outputs to decrease runtime. The Galaxy integration of EDTA and panEDTA is accessible through: <<https://github.com/bgruening/galaxytools/pull/1244>>, which can be deployed in local servers following the general Galaxy guideline (https://galaxyproject.org/admin/get-galaxy/).

### Genomes and TE annotations

Fifty Arabidopsis genomes were downloaded from NCBI (**Suppl. Data 1**) and annotated by panEDTA. The curated Arabidopsis library was obtained from RepBase 20.03 (ref ^46^) and reformatted to follow the naming conventions of panEDTA. The curated Arabidopsis library was provided to panEDTA with the “--curatedlib” parameter.

This study utilized previously generated genome assemblies of the 26 maize NAM founder lines^17^, which included 13 tropical genomes (CML103, CML228, CML247, CML277, CML322, CML333, CML52, CML69, Ki3, Ki11, NC350, NC358, and Tzi8), 10 temperate genomes (dent genomes: B73, B97, Ky21, M162W, Ms71, Oh43, and Oh7B; flint genomes: HP301, P39, and Il14H), and three admixed genomes (M37W, Mo18W, and Tx303). Briefly, these genomes were sequenced to high depth (63-85x) using the PacBio CLR technique and assembled to high contiguity with an average contig N50 of 25.7 Mb, including complete assembly of the TE space based on the LTR Assembly Index (LAI)^18^ with a mean value of 28 (ref ^17^). In addition, the *Zea mays* ssp. *mexicana*, PI 566673 genome (GenBank PRJNA299874) was used as an outgroup to polarize LTR insertions, which was sequenced using PacBio CLR and Illumina HiSeq 2000 (ref ^47^).

Although the lines used to generate the 26 maize NAM founder genomes are highly inbred, there are still low levels of heterozygosity present as alternative scaffolds in these genomes. We collected known alternative scaffolds from MaizeGDB (https://www.maizegdb.org/), and further identified the remaining alternative scaffolds based on alignments to pseudomolecules. Unplaced scaffolds in each genome were aligned to their respective pseudomolecules using minimap2 (v2.24)^48^, and those having a mapping quality of 60 were considered alternative scaffolds (-minq 60). To avoid over-counting of haploid assembly size and transposable element content, alternative scaffolds collected from MaizeGDB and identified in this study were removed from downstream analyses.

Using panEDTA, we generated a new version of the TE annotation for the 26 maize NAM founder genomes. Names of sequences in the panEDTA library were ported from the original MTEC library names^2^ or generated by EDTA for novel TEs in the panEDTA library that were not previously contained in the MTEC library. Annotation and classification of TE families are available from MaizeGDB: https://ars-usda.app.box.com/v/maizegdb-public/folder/176298403241 (“EDTA.TEanno.gff3” files). The size of each TE family in bp was summarized from the annotation of each genome using the “buildSummary.pl” script derived from the RepeatMasker package^49^.

### Annotation evaluation

Repeat annotations (RepeatMasker “.out” files) generated using EDTA and panEDTA libraries were evaluated for annotation inconsistency by the script “evaluation.pl” in the EDTA package. The maize B73v5, rice MSU7, and Arabidopsis TAIR10 genomes were used for the evaluation. The rice EDTA and panEDTA library was obtained from ref ^50^, and the maize and the Arabidopsis EDTA and panEDTA libraries were generated in this study. Such evaluations are reference-free and thus do not rely on the availability of a gold-standard annotation. Briefly, annotated repeat sequences of a genome were extracted and subjected to all-vs-all blast within the extracted sequences, and the matching sequences covering ≥95% of the query with ≥80% identity and ≥80 bp in length were compared to the query sequence’s annotation to identify inconsistently annotated entries. The annotation inconsistency was measured at the superfamily level.

### Pan-genome analysis

To estimate the distribution of TE families in the pan-NAM founder genomes, only pan-genome families that contained at least one full-length TE (fl-TE) in at least one of the genomes were included. Full-length TEs were identified using the pan-genome TE annotation and the “find_flTE.pl” script in the EDTA package. The lists of fl-TE families from the NAM founder genomes were added incrementally (from 1 to 26 genomes) and in random order. The number of unique pan-genome fl-TE families was counted after adding each genome’s fl-TE list. This process was iterated 1,000 times.

Principal component analysis (PCA) of pan-NAM TE families (n = 17,473) was computed in R version 4.0.3 (ref ^51^) using the command “prcomp” and the physical size of each family in bp in each genome. TE family sizes were assigned as 0 in the genomes that are absent. Unnormalized family sizes were used so that larger TE families will have more weight in the PCA. The SNP PCA was done using 25,000 random homozygous biallelic SNPs with no missing data that were filtered from the original NAM SNP dataset^17^ (https://datacommons.cyverse.org/browse/iplant/home/shared/NAM/NAM_genome_and_annotation_Jan2021_release/SUPPLEMENTAL_DATA/NAM-founder-SNPs) in R using the command “prcomp.” The reference and alternative alleles were coded as 0 and 1, respectively. Unnormalized values were also used to conduct the SNP PCA.

To construct the phylogeny of the 26 NAM founder genomes, BUSCO genes were identified in all of the 26 NAM founder genomes and *Zea mays* ssp. *parviglumis* using BUSCO (v3.1.0)^52^ with the Embryophyta odb9 dataset (n = 1,440) and the Augustus species ‘maize’ parameters. Subsequently, amino acid sequences of the complete BUSCO genes were extracted and subjected to multiple sequence alignment using GUIDANCE (v2.0)^53^ with “mafft” selected as the alignment program. GUIDANCE provides a framework for identifying and removing phylogenetically unreliable regions in the multiple sequence alignment. This multiple sequence alignment was then used to create the phylogenetic gene tree using the maximum likelihood approach in RAxML (v8.2.11)^54^, with *Zea mays* ssp. *parviglumis* as an outgroup. RAxML was run with 100 bootstrap replicates with the option PROTGAMMAAUTO to automatically determine the best-fitting protein substitution model for the data. Trees obtained from the different bootstrap runs were then merged to generate a consensus gene tree. This consensus gene tree was used as an input to generate the final species tree for the NAM genomes using ASTRAL (v4.10.7)^55^ with default parameters. ASTRAL is a fast and accurate method for estimating a species tree from gene trees based on a multispecies coalescent model. The resulting species tree was plotted using ggtree (v2.4.1)^56^.

### Copy number estimates for unclassified LTR retrotransposons

Some intact LTR retrotransposons (LTR-RTs) could not be classified into families due to low copy numbers in the original TE annotation and were thus named by their coordinates in the genome. To estimate the copy number of these unclassified LTR-RTs, their sequences were extracted in each genome and redundant copies were removed using the “cleanup_nested.pl” script from the EDTA package with parameters ‘-cov 0.95 -minlen 80 -miniden 80’ that require at least 80 bp, 80% identity, and 95% sequence coverage. The resulting representative sequences were used to mask the original LTR sequence using RepeatMasker with parameters ‘-q -no_is - norna -nolow -div 40’ that allows for up to 40% of divergence. Masked sequences were annotated by the “classify_by_lib_RM.pl” script using relaxed parameters ‘-cov 50 -len 70 -iden 70’ that require at least 70 bp, 70% identity, and 50% coverage. After this re-annotation step, the copy number of representative intact LTR-RTs was counted in the re-annotated sequences.

### Metadata of intact LTR retrotransposons

The length, classification, and divergence information of each intact LTR-RT were obtained from the original annotation of each genome (“pass.list” files)^17^. The insertion time of each intact LTR element was calculated from the LTR divergence using the maize molecular clock μ = 3.3e-8 per bp per year^57^. For each intact LTR-RT, nested TEs were identified when other TEs were fully enclosed in the intact LTR-RT, and the copy number and length of the nested TEs were counted for each intact LTR-RT. The number of conserved coding domains in each intact LTR-RT was identified using TEsorter (v1.3)^58^ with default parameters. Gene annotations used in this study were filtered from the original NAM gene dataset^17^ (https://datacommons.cyverse.org/browse/iplant/home/shared/NAM/NAM_genome_and_annotation_Jan2021_release/GENE_MODEL_ANNOTATIONS), in which only primary isoforms were retained (“T001” transcripts). TE-related genes were identified using TEsorter with default parameters and were removed. The physical distance of each intact LTR-RT to the downstream gene is calculated by “closest-features --dist” function with the BEDOPS (v2.4.39) package^59^. Solo LTRs were identified using the “solo_finder.pl” script from the LTR_retriever package^23^ and modified to adapt to the current annotation format. For each family in each genome, the solo:intact ratio was calculated using the number of solo LTRs over the number of intact LTR-RTs for the LTR family. Data of all solo and intact LTRs in each genome can be accessed from MaizeGDB: https://ars-usda.app.box.com/v/maizegdb-public/folder/176298403241.

### Classification of LTR family dynamics

Family size and solo:intact ratio of each family was compared between tropical genomes (n = 13) and temperate genomes (n = 10) to determine LTR family dynamics. Comparisons between tropical and temperate genome groups were based on a Student’s *t-*test and *P* ≤ 0.05 as the significance cutoff. Families with significantly different family size and solo:intact ratio were classified as removal families. Families with significantly different family size but no statistical difference in solo:intact ratio were classified as amplification families. Families with significantly different solo:intact ratio but no statistical difference in family size were classified as balanced families. Finally, families with neither significantly different family size nor solo:intact ratio were classified as drifting families. For amplification families, if the average family size was larger in tropical than in temperate genomes, the family was classified as a tropical amplification family, and conversely, if that in the temperate genomes were larger the family was classified as a temperate amplification family. Likewise, for removal families, if the average family size was smaller in tropical than in temperate genomes, the family was classified as a tropical removal family, and conversely, if the average family size was smaller in the temperate genomes the family was classified as a temperate removal family. A small number of removal families have inconsistent direction of removal and family size between tropical and temperate genomes, specifically, tropical removal families having solo:intact ratio higher in temperate genomes and temperate removal families having solo:intact ratio higher in tropical genomes. These families contributed only small effects to TE content variation (-0.20 Mb and 0.59 Mb, respectively), and were removed from downstream analysis. All family classifications can be found in **Suppl. Data 2**).

### Classification of LTR families based on Age

LTR families were classified into Young, Moderate, and Old using the age distribution of intact LTR-RTs in each family within the temperate and tropical genomes (**Suppl. Table 1**). LTR families with less than 10 copies were not classified due to low confidence in inferring their age distributions. For those families with greater than 10 copies among the temperate and tropical genomes, the divergence of intact LTR-RTs was binned with 0.002 identity intervals and the frequency of intact LTR-RTs was calculated in each bin. If the first bin ([0, 0.002)) had the highest frequency, the family was classified as a “Young” family. If the first bin did not have the highest frequency but contained ≥5% total LTR-RTs of this family, the family was classified as a “Moderate” family. If the first bin contained <5% total LTR-RTs of this family, the family was classified as an “Old” family.

### Determining the epigenetic status of LTR elements

Unmethylated regions (UMRs) were previously identified based on enzymatic methyl-seq reads (PE 150, ≥300M reads per genotype) from second leaves of pooled plants with two biological replications for each genome^17^. These UMRs were defined primarily based on hypomethylation in the CHG context (H = A, T, or C), which is a strong indicator of euchromatin. Many of these CHG-defined UMRs contain high levels of methylation in the CG context^17^. Only intact LTR elements with unambiguous strand directions were used for this analysis. The coordinate of the 5’ LTR of each element was determined based on the pan-genome annotation and the strand information, which was used to intersect with whole-genome UMRs using BEDTools (v2.30.0)^60^. UMRs that overlapped ≥ 200 bp with the 5’ LTRs were candidates of unmethylated LTRs. Those that started upstream of the 5’ LTR on the correct strand were removed. The remaining UMRs were determined to have originated within 5’ LTRs (UM-5’LTRs).

Sequences for UMRs originating within the centromeric retrotransposons of maize 2 (CRM2) elements were extracted from NAM founder lines, and only those located on the positive strand were retained. MAFFT (v7.487) was used to align CRM2-UMRs to the CRM2 sequence from the TE library with default parameters. The resulting alignment was converted to the SAM format using JVarkit biostar139647 (ref ^61^) and visualized using IGV (v2.4.17)^62^.

### TE family expression analysis

Family level transcript abundance estimates were computed for two replicates of 10 tissues for each genotype^17^ using a previously described method^32^ adapted for NAM TE annotations. Briefly, reads were mapped using HiSat2 (v2.1.0) (parameters -p 6 -k 20)^63^, sorted by name using samtools sort (v1.9)^64^, and overlapped with features using HTseq (parameters -s no -t all - m union -a - --nonunique all). The annotation files used in HTseq were generated by first subtracting exon regions from the TE annotation for each NAM genome using BEDTools subtract (v2.27.1)^60^, then concatenating this file with the full-length gene sequences before sorting. This resulting annotation file prioritizes genes over TEs in overlapping regions. Count tables for each TE family and all genes (collapsed into an entry named “Gene”) were then created with the script “te_family_mapping_ver8.2_NAM.pl.” This script counts reads towards TE families if they are uniquely mapping to a single TE or the read is multi-mapping and all mapping locations that intersect a feature are annotated as the same TE family. Paired-end reads were only counted once.

The table containing raw read counts for each TE family (**Suppl. Data 3**) was used to identify differentially expressed families between tropical and temperate genomes. Only families that were shared by tropical and temperate genomes were retained (n = 15,957). Libraries from the three admixed genomes (M37W, Mo18W, and Tx303) were removed. To normalize the library size effect caused by differences in sequencing depth, total read counts for TEs were considered, and the “median of ratios” method^65^ was used to estimate the normalization factor for each library (**Suppl. Data 4**). In brief, for each library, raw counts were divided by the geometric mean of each TE family. The median for non-zero ratios in a library was used as the size factor for this library. The variance stabilizing transformation (VST) was then used to transform and normalize raw counts using DESeq2 (v1.30.1)^66^ based on the normalization factors determined using the median of ratios method. TE families differentially expressed between tropical and temperate genomes were identified for each of the 10 tissues and for all tissues with adjusted *P* values < 0.05 (Wald test, FDR) using DESeq2 (v1.30.1)^66^. To normalize for family size, FPKM (fragments per kilobase of sequence per million mapped fragments) values of each family were estimated using all tissues and replicates combined. The total length of each TE family was used in the FPKM calculation, and the log2 + 1 method was used to transform raw FPKM values.

### Identification of syntenic LTR retrotransposons

Syntenic intact LTR-RTs^67^ were identified from the panEDTA annotation of the intact LTR-RTs based on syntenic information flanking each element. Pairwise syntenic LTR-RTs were identified in pairs of two genomes with a total of 325 combinations of pairs between the 26 NAM founder genomes. To identify syntenic LTR-RTs between two genomes, two 1 kb sequences centered on the start and end position of each intact LTR-RT in both genomes were extracted and blasted against the paired genome. Blast hits to the orthologous chromosome (e.g., an intact LTR-RT from B73 Chr1 hitting the Chr1 of Tzi8) with an e-value < 1e-5 and query coverage > 40% were retained as candidate hits. Each hit was further classified as a full match if there was > 75% query coverage or a half match if there was ≤ 75% query coverage. Half matches were discarded if the majority of the hit (> 50% query coverage) was from the LTR portion of the 1-kb sequence, as these hits have a high probability of being off-target matches. If full matches were found for both 1-kb sequences from the LTR-RT in the same direction and the hits were ≥ 200 bp apart (non-empty sites) and less than or equal to the length of the intact LTR-RT plus 100 kb, then the locus was recorded as the syntenic full site of the intact LTR-RT.

These parameters allow for both syntenic intact LTR-RTs and syntenic solo LTR-RTs to be identified (**Suppl. Fig. S16**). The lack of an identified insertion could be due to the full deletion of a preexisting LTR-RT insertion or the lack of insertion (a null site). To identify null sites, we required that half matches were found for flanking sequences of both the 1-kb sequences and that these were ≤ 10 bp apart. If an LTR-RT was not identified as a full site or a null site in the paired genome based on these criteria, it was recorded as missing data. If more than one full site or null site on the orthologous chromosome were observed, the LTR-RT was considered recalcitrant within a particular pair of genomes and was recorded as missing data in the paired genome. After the identification of syntenic loci in a genome, their coordinates were used to overlap with the TE annotation of that genome to obtain the exact coordinate of the syntenic LTR-RT. Because syntenic LTR-RTs were identified using all intact LTR-RTs of the genome pair, we further separated queries back to two genomes, resulting in reciprocal pairwise syntenic LTR-RT information with a total of 650 pairs of genomes.

To create the pan-LTR matrix, reciprocal pairwise syntenic LTR-RT results were joined in R (v3.6.3) using the left_join() function for each genome respectively via the script “create_pan_matrix_by_genome.R.” The LTR matrix for the 26 genomes was concatenated, then duplicated LTR coordinates were compressed using the script “compressing_duplicate_TE.R”. Briefly, LTR coordinates within each genome were searched for duplicates. Duplicate coordinates were compressed and LTR coordinates in other genomes were merged. For LTRs that have more than one hit after the compression, the LTRs were stored as semicolon-separated pairs. Finally, the “Resolve_conflict.py” script was used to merge LTRs coordinates. For the semicolon-separated LTRs, the entry containing an intact LTR was prioritized to retain, followed by truncated and null sites. LTRs whose presence/absence status could not be determined based on these parameters were considered missing data and demarcated as NA in the matrix. Data from the three admixed genomes (M37W, Mo18W, and Tx303) were removed from this study but retained in the data repository. Syntenic LTR-RTs with missing data rates over 50% within the tropical genomes, within the temperate genomes, or across all genomes were discarded.

To obtain the ancestral state of syntenic LTR-RTs, we extracted 500 bp sequences flanking the insertion site of syntenic LTR-RTs, with 250 bp on each side. Flanking sequences were combined to form 500 bp null site sequences and used to blast against the *Zea mays* ssp. *mexicana* genome. Null sites in the *Zea mays* ssp. *mexicana* genome were identified when a single blast hit covering ≥80% of the query with ≥95% identity was found in the genome. An age cutoff of ≤20 kya was also used to identify syntenic LTR-RTs that were inserted after the divergence between cultivated maize and its wild progenitor teosinte^68^ and compared to the *Zea mays* ssp. *mexicana-*based polarization. Data of all synLTRs in each genome can be accessed from MazieGDB: https://ars-usda.app.box.com/v/maizegdb-public/folder/176298403241.

LTR insertion frequency spectrums were estimated on both missing-filtered and missing-unfiltered datasets using SoFoS (v2.0) (https://github.com/CartwrightLab/SoFoS) with parameters “-r -a 1.0 -b 1.0.” The population size was rescaled to 10 (-n 10) to account for imbalanced population size between tropical and temperate groups with either folded (-f) and unfolded (-u) estimations.

### Estimation of recombination rate

A composite recombination map derived from all NAM recombinant inbred lines (RILs, backcrossed to B73) across all families was obtained from ref ^35^ and used to estimate the local recombination rate at each of the intact LTR-RTs. The genetic map was downloaded from the CyVerse data commons (https://datacommons.cyverse.org/browse/iplant/home/silastittes/parv_local_data/map/ogut_v5.map.txt), and converted to recombination rate in the unit of cM/Mb in R (v4.0.3)^51^ based on the B73v5 physical map. The recombination rate at each intact LTR-RT was approximated using the recombination rate data point with the nearest physical distance.

### Estimation of allele frequency using SNPs

To call SNPs for NAM founders against the *Zea mays* ssp. *mexicana* genome (GenBank PRJNA299874), we used the Genome Analysis Toolkit (GATK v4.2.2.0) and followed the Bioinformatics Workbook (https://bioinformaticsworkbook.org/dataAnalysis/VariantCalling/gatk-dnaseq-best-practices-workflow.html). In brief, short reads from the 26 NAM genomes were downloaded from CyVerse (ENA PRJEB31061) and were used for calling SNPs by mapping to the *Zea mays* ssp. *mexicana* genome as the reference. The GATK HaplotypeCaller^69,70^ and the Picard Toolkit (v2.26.6) (http://broadinstitute.github.io/picard/) were used for SNP discovery and final variant filtering. Picard FastqToSam was used to convert fastq format to SAM format, then Picard MarkIlluminaAdapters was used to mark Illumina adapters and generate metrics files. The SAM formatted files were converted back to interleaved fastq files using the Picard SamToFastq utility, which were then mapped to the BWA-MEM-indexed *Zea mays* ssp. *mexicana* genome using recommended options (-M)^71^. The aligned reads were merged with unaligned reads using Picard MergeBamAlignment, marking duplicates with Picard MarkDuplicates. In the last step of processing BAM files, AddOrReplaceReadGroups was used to add the correct read-group identifier before calling variants with HaplotypeCaller. HaplotypeCaller was trivially parallelized by running simultaneously on 2-Mb intervals of the genome (587 chunks, excluding scaffolds), and the VCF files were gathered to generate a merged, coordinate-sorted, unfiltered set of SNPs. Stringent filtering was performed on the raw set of SNPs using the expression (QD < 2.0 || FS > 60.0 || MQ < 45.0 || MQRankSum < -12.5 || ReadPosRankSum < -8.0 || DP > 5061), where DP was estimated from the DP values of the SNPs (standard deviation times 5 + mean). This set was filtered to retain only homozygous, biallelic SNPs, which are available via MaizeGDB: https://ars-usda.app.box.com/v/maizegdb-public/folder/176298403241. BCFtools (v1.9)^72^ was used to control missing data rate ≤50% with parameters “view -e ‘F_MISSING>=0.5’.” The VCFtools (v0.1.16)^73^ vcf-subset utility was used to split VCF files into tropical and temperate subpopulations. Folded SNP SFS were estimated on both missing-filtered and missing-unfiltered SNP sets for both tropical and temperate subpopulations using SoFoS with parameters “-f -r -a 1.0 -b 1.0” and population sizes rescaled to 10 (-n 10).

### Statistical analyses

All statistical analyses and graphic visualizations in this paper were performed using R (v4.0.3)^51^ in RStudio (v1.1.442)^74^. Aesthetic modification and compilation of plots were done using Inkscape (v1.0) (https://inkscape.org).

## Data and Code availability

Arabidopsis genomes were downloaded from NCBI (**Suppl. Data 1**). Maize genome assemblies, gene annotations, and pan-genome TE annotations were downloaded from https://maizegdb.org/NAM_project. Illumina resequencing reads were downloaded from ENA PRJEB31061. RNA-seq reads were downloaded from ENA ArrayExpress E-MTAB-8633 and E-MTAB-8628. Enzymatic methyl sequencing reads were downloaded from ENA ArrayExpress E-MTAB-10088. Scripts and files used to generate and analyze data are available on GitHub: https://github.com/oushujun/PopTEvo and MaizeGDB: https://ars-usda.app.box.com/v/maizegdb-public/folder/176298403241.

## Acknowledgments

We thank the Zeavolution online seminar series for providing critical feedback and suggestions based on an earlier version of this study. We thank Dr. Guanjing Hu for helpful guidance and discussion regarding analyses of expression data. We thank Alex Ostrovsky, Björn Grüning, and the entire Galaxy community for assistance in integrating panEDTA. We thank MaizeGDB for hosting the data generated in this study. This work was supported in part by the National Science Foundation (Grants IOS-1546727, IOS-1934384, IOS-1744001, IOS-1758800, IOS-2216612, and IOS-1546719), National Institutes of Health (U24HG006620), the Human Frontier Science Program (RGP0025/2021), the Minnesota Agricultural Experiment Station, and the Iowa State University Postdoctoral Seed Grant (PG101847). The computation was performed in part using the HPC platform at Iowa State University, which was purchased in part by the NSF grant MRI-1726447, and in part using the Minnesota Supercomputing Institute (MSI) at the University of Minnesota.

## Author information

### Affiliations

**Department of Ecology, Evolution, and Organismal Biology, Iowa State University, Ames, USA**

Shujun Ou, Arun S. Seetharam, Nancy Manchanda, Matthew B. Hufford

**Department of Agronomy and Plant Genetics, University of Minnesota, St. Paul, USA**

Shujun Ou, Yinjie Qiu, Claire C. Menard, Candice N. Hirsch

**Department of Computer Science, Johns Hopkins University, Baltimore, MD, USA**

Shujun Ou, Tyler Collins, Michael C. Schatz

**Department of Genetics, Development, and Cell Biology, Iowa State University, Ames, USA**

Sarah N. Anderson, Arun S. Seetharam

**Department of Plant Biology, University of Georgia, Athens, USA**

Jonathan I. Gent

### Contributions

S.O., A.S.S., N.M., Y.Q., and S.N.A. generated and curated the data. N.M. performed the phylogenetic analysis. Y.Q. combined pairwise syntenic LTR information into the population level. S.N.H. generated the family-based TE expression data and assisted in analyzing them. J.I.G. assisted in planning and interpreting DNA methylation analyses. A.S.S. performed read mapping, SNP calling, and assisted in recombination analyses. S.O. performed the pan-genome TE annotations, curations, and summaries, classified and analyzed LTR family age and dynamic groups, identified and analyzed UM-5’LTRs, analyzed family-based TE expression data, identified and analyzed syntenic LTR-RTs, analyzed recombination data, and estimated SFS. C.C.M. performed quality control analyses of the TE annotations. T.C. implemented the Galaxy instance of the panEDTA module. S.O., T.C., A.S.S., N.M., Y.Q., M.B.H., and C.N.H. prepared the manuscript and all authors revised it. M.C.S., M.B.H., and C.N.H. are senior authors who secured funding and oversaw the project.

### Corresponding authors

Correspondence to Matthew B. Hufford or Candice N. Hirsch.

